# Was the extinct Spectacled Cormorant part of the North Pacific kelp-associated fauna?

**DOI:** 10.64898/2026.09.02.749039

**Authors:** Amy L. Adams, Pascale Lubbe, Alexander T. Salis, Martync Kennedy, Robert Boessenecker, Alexander L. Bond, Hamish G. Spencer, Nicolas J. Rawlence

## Abstract

The coastal North Pacific kelp ecosystems are highly productive, supporting a rich faunal assemblage. Some historical aspects of this ecosystem remain poorly studied, as exemplified by the extinct Spectacled Cormorant *Urile perspicillatus*, whose Late Holocene distribution was apparently restricted to Bering Island. This cormorant’s taxonomic relationships remain unresolved and little is known about its ecology. We use ancient-DNA to reconstruct the evolutionary history of the Spectacled Cormorant. Mitogenomes from five of the six known historical skins form a strongly supported clade within *Urile* cormorants. The ancestors of Spectacled Cormorant diverged from its closest relatives, the Pelagic (*U. pelagicus*) and Red-faced (*U. urile*) cormorants, ∼6.98 Mya, and the population subsequently underwent a pronounced bottleneck. This divergence time is co-incident with the establishment of coastal kelp-forests. The Spectacled Cormorant may have been an integral part of the kelp-associated fauna as a large benthic-foraging predator. Similar to the sympatric Steller’s sea cow (*Hydrodamalis gigas*), the historical relict distribution of the Spectacled Cormorant may be the result of trophic collapse of kelp-forest ecosystems and subsistence hunting. Future palaeogenomic and dietary isotopic research, and reassessment of North Pacific archaeological and fossil assemblages, should shed light on the evolution and palaeoecology of this enigmatic seabird.

## INTRODUCTION

The kelp forests of the North Pacific are a highly productive coastal marine ecosystem that provides important habitat for a diverse associated fauna (Estes *et al*. 2016; Vermeij *et al*. 2019; Kiel *et al*. 2024). Although research is beginning to shed light on the evolution and development of this rich ecosystem and how it has changed through time (e.g. Estes *et al*. 2016; Starko *et al*. 2019), there are several aspects of its character that remain enigmatic and poorly known. This situation is exemplified by the recently extinct Spectacled Cormorant *Urile perspicillatus* (Pallas, 1811), whose ecology is virtually unknown and limited to observations by the naturalist Georg Wilhelm Steller.

*Urile perspicillatus* may be the largest phalacrocoracid to have ever lived, with some sources suggesting it weighed up to 6.8 kg (Hume 2017) and stood ∼100 cm in height (Johnsgard 1993; Figure 1C). At the time of European discovery in 1741, the species was “exceedingly common” (based on Stejneger’s [1936] translation of the Latin “copiosissimi” in Steller’s unpublished notes) but restricted to Bering Island in the Commander Islands between the Kamchatka Peninsula and the Aleutian arc (Figure 1A). It may also have been found on nearby islands (Pallas 1811) and as a rare vagrant to Kamchatka (Fuller 2000). Traditional ecological knowledge suggests that *U. perspicillatus* may have also inhabited the Near Islands >300 km to the east of Bering Island (Turner 1886), though specimens of Holocene age from anywhere but Bering Island have yet to be described (Squires & Bond 2024; Samsonov *et al*. 2026). Like the more well-known sympatric Steller’s sea cow *Hydrodamalis gigas* (Zimmermann, 1780) (Turvey & Risley 2006), *U. perspicillatus* was driven to extinction through over-harvesting within one hundred years of European discovery, though predation by Arctic foxes *Vulpes lagopus* (Linnaeus, 1758), volcanic activity, and disease could have also contributed (Stejneger 1883; Stejneger 1885; Stejneger and Lucas 1889; Hartert 1920; Stejneger 1936; Greenway 1967; Johnsgard 1993; Fuller 2000; Hume 2017).

**Figure 1.**
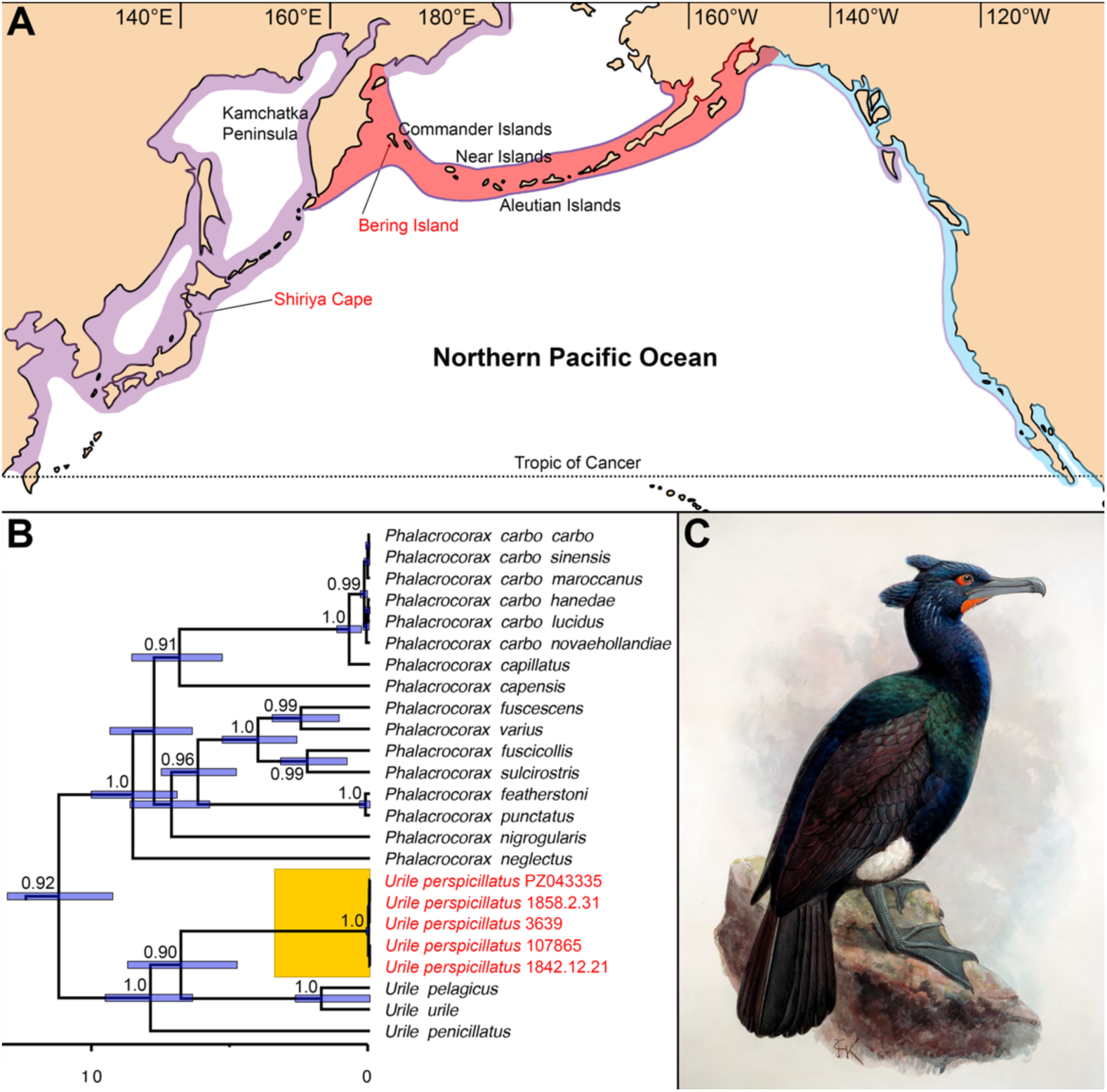
**A**: Stylized distribution of *Urile* cormorants based on species accounts from Birds of the World (Causey 2021; Hobson 2021; Wallace & Wallace 2021). Purple: Pelagic Cormorant (*U. pelagicus*); red: Red-faced Cormorant (*U. urile*); blue: Brandt’s Cormorant (*U. penicillatus*). The purple outline on the distribution of *U. urile* and *U. penicillatus* indicates the sympatric distribution of *U. pelagicus*. Late Holocene and Late Pleistocene specimens of Spectacled Cormorant (*U. perspicillatus*) have been found on Bering Island and Shiriya Cape, respectively. **B**: Evolutionary history of *U. perspicillatus* and its nearest relatives. The time-calibrated phylogeny (in millions of years) is derived from 3.5 kilobases of mitochondrial DNA sequence data; the StarBEAST species-tree analysis additionally incorporated five nuclear loci for comparative taxa. Node bars on the phylogeny are 95% HPD of divergence times as indicated on the scale bar (millions of years before present). Node values are shown for BEAST Bayesian posterior probability supports of 0.90 and above. For the full phylogeny see Supplementary Figure S2. **C**: Artistic reconstruction of *U. perspicillatus* by J. G. Keulemans (Image courtesy of the Natural History Museum, Tring).

Ecological knowledge of *U. perspicillatus* comes exclusively from Steller’s observations and fragmentary Indigenous sources (Pallas 1811; Stejneger 1883; Stejneger 1936; Turner 1886). It has been described as large, clumsy, slow on land, and flight-reduced (Domning 1978; Fuller 2000; Hume 2017). However, Samsonov *et al*. (2026) hypothesized that this taxon was not flight reduced, merely reluctant to fly, based on morphometric analysis of a large collection of late Holocene fossil bones from Bering Island (supporting previous studies; i.e., Palmgren 1935; Livezey 1992). Samsonov *et al*. (2026) suggested its large size (and hindlimb morphology) may have evolved in response to deep diving for benthic prey, similar to blue-eyed shags (*Leucocarbo* spp.), which are poor fliers (Rawlence *et al*. 2022). Furthermore, weight calculations by Samsonov *et al*. (2026) suggest *U. perspicillatus* weighed only 2.5-4.5 kg (in contrast to Steller’s recorded 12-14 pounds; Stejneger and Lucas 1889). It should be noted that equations to calculate body mass from bone measurements can be problematic (e.g. Rawlence *et al*. 2025), especially when compared to live caught birds in which weight can fluctuate seasonally and with feeding events (Koffijberg & van Eerden 1995). In addition, an 18^th^ Century Russian pound is c. 409 g (*cf*. an imperial pound at 450 g), which would imply a weight range for *U. perspicillatus* of 4.9-5.7 kg (see Samsonov *et al*. 2026), compared with, for example, the 5.5-6.35 kg range estimated in Johnsgard (1993) using the imperial pound weight.

There are currently six known historical skins of *U. perspicillatus*, but hundreds of historic and fossil bones (mostly Late Holocene in age), as detailed by Squires and Bond (2024) and Samsonov *et al*. (2026). Plumage analysis has traditionally suggested that the Spectacled Cormorant (Figure 1C) is closely related to those species in the genus *Urile* that inhabit the northern Pacific (Figure 1A): Brandt’s Cormorant *U. penicillatus* (Brandt, 1837), found along the west coast of North America; Red-faced Cormorant *U. urile* (Gmelin, JF, 1789), which occurs along the Aleutian Island chain from Alaska to the Kamchatka Peninsula; and the Pelagic Cormorant *U. pelagicus* (Pallas, 1811), found in coastal regions from Baja California to Japan. Based on the osteological analysis of Holocene fossil bones, it has been hypothesized that *U. perspicillatus* is most closely related to *U. urile* and *U. pelagicus* (Samsonov *et al*. 2026).

The recent discovery of ∼120,000-year-old fossil bones of *U. perspicillatus* from Shiriya Cape in northeastern Honshu (Aomori Prefecture), Japan (Figure 1A) points to the possibility of the historical range being a relict of a formerly more widespread Pleistocene distribution or a population shift due to changing ocean productivity (Watanabe *et al*. 2018). A dramatic reduction in range and/or population size in response to changing conditions after the Last Glacial Maximum may have resulted in increased extinction risk in the face of significant anthropogenic pressure. This significant fossil find also raises the possibility of a closer phylogenetic affinity of *U. perspicillatus* to large cormorants from Japan, specifically the Great Cormorant *Phalacrocorax carbo* (Linnaeus, 1758) and Japanese Cormorant *P. capillatus* (Temminck & Schlegel, 1850) (Rawlence *et al*. in review), both of which share some plumage similarities with *U. perspicillatus*.

Here we sequence mitogenomes from four of the six known historical specimens of *U. perspicillatus*. We use these data to estimate a molecular-dated phylogeny to reconstruct the evolutionary history of this species to (1) test whether *U. perspicillatus* is a *Phalacrocorax* or *Urile* cormorant; (2) determine when this large cormorant evolved and what may have driven its evolutionary history; and (3) investigate the presence of a population bottleneck associated with prehistoric range contraction and/or recent human over-exploitation.

## MATERIALS AND METHODS

### Sampling and laboratory protocols

Toe pads were sampled from four of the six known historical skins of *U. perspicillatus*: NHMUK 1842.12.21.4 and 1858.2.3.1 (Natural History Museum, Tring), RMNH.AVES.107865 (Naturalis Biodiversity Center, Leiden), and MZH UL 3639 (Luonnontieteellinen keskusmuseo, LUOMUS - Finnish Museum of Natural History, Helsinki). Permission to sample the two specimens in the Zoological Institute of the Russian Academy of Sciences in St. Petersburg, Russia, was not forthcoming. However, a fifth mitogenome presumably from one of these historical skins was recently released on GenBank (PZ043335.1). None of the skins have a known provenance as all were routed through present-day Sitka, Alaska, the former capital of Russian America, in the 1830s and 1840s (Squires & Bond 2024).

Ancient DNA extraction and library preparation were performed in a dedicated, physically isolated, ancient DNA laboratory (Ancient Ecology Lab) following strict ancient DNA guidelines, including the use of negative extraction and no-template library controls (Knapp *et al*. 2012). Extractions followed a modified protocol in which toe pads were agitated for 30 seconds in 100 µL 0.5 M EDTA before being minced and incubated overnight at 50°C in a rotator in 180 µL Buffer ATL (Qiagen), 20 µL 10 mg/mL proteinase K, and 20 µL 1 M DTT following Fulton *et al*. (2011). Ancient DNA extracts were purified and eluted in 50 µL TET buffer following Dabney *et al*. (2013).

Single-stranded DNA libraries were constructed following Gansauge *et al*. (2017). The optimal number of indexing Polymerase Chain Reaction (PCR) cycles was determined by quantitative PCR, identified as the point at which the amplification curve plateaued. Indexed libraries were purified using the MinElute PCR purification kit (Qiagen). Library concentrations were quantified using a Qubit fluorometer (Thermo Fisher Scientific), while fragment length distributions were determined using a TapeStation (Agilent) to calculate library molarity. Libraries were pooled and sequenced with 50 base pair (bp) paired-end reads (2x50 bp) on an Illumina NextSeq 2000 sequencer at the Otago Genomics sequencing facility.

### Bioinformatic and phylogenetic analyses

Illumina adapter sequences and low-quality bases were trimmed from de-multiplexed sequencing reads using fastp v0.23.4 (Chen *et al*. 2018), applying a minimum quality threshold (-q) of 30 and a minimum read length (-l) of 25, following which reads were collapsed based on a minimum overlap length (--overlap_len_require) of 20. To identify the most appropriate reference scaffold, collapsed reads were mapped to publicly available reference mitogenomes from several closely related species: *P. capillatus*; *P. carbo carbo* (Linnaeus, 1758); *P. carbo hanedae* Kuroda, 1925; *U. perspicillatus*; and *U. pelagicus*; Table 1) using BWA aln (Li & Durbin 2009) with parameters recommended for degraded DNA (Schubert *et al*. 2012): a maximum edit distance of 0.01 (-n), a maximum of 2 gaps open (-o), and seeding disabled (-l 1024). Alignments with mapping quality (-q) below 30 were removed using Samtools v1.6 (Li *et al*. 2009), and PCR duplicates were removed using the MarkDuplicates function in Picard v2.23.4 (Broad Institute 2019). Summary mapping statistics were generated using SAMtools flagstat (Li *et al*. 2009).

**Table 1.**
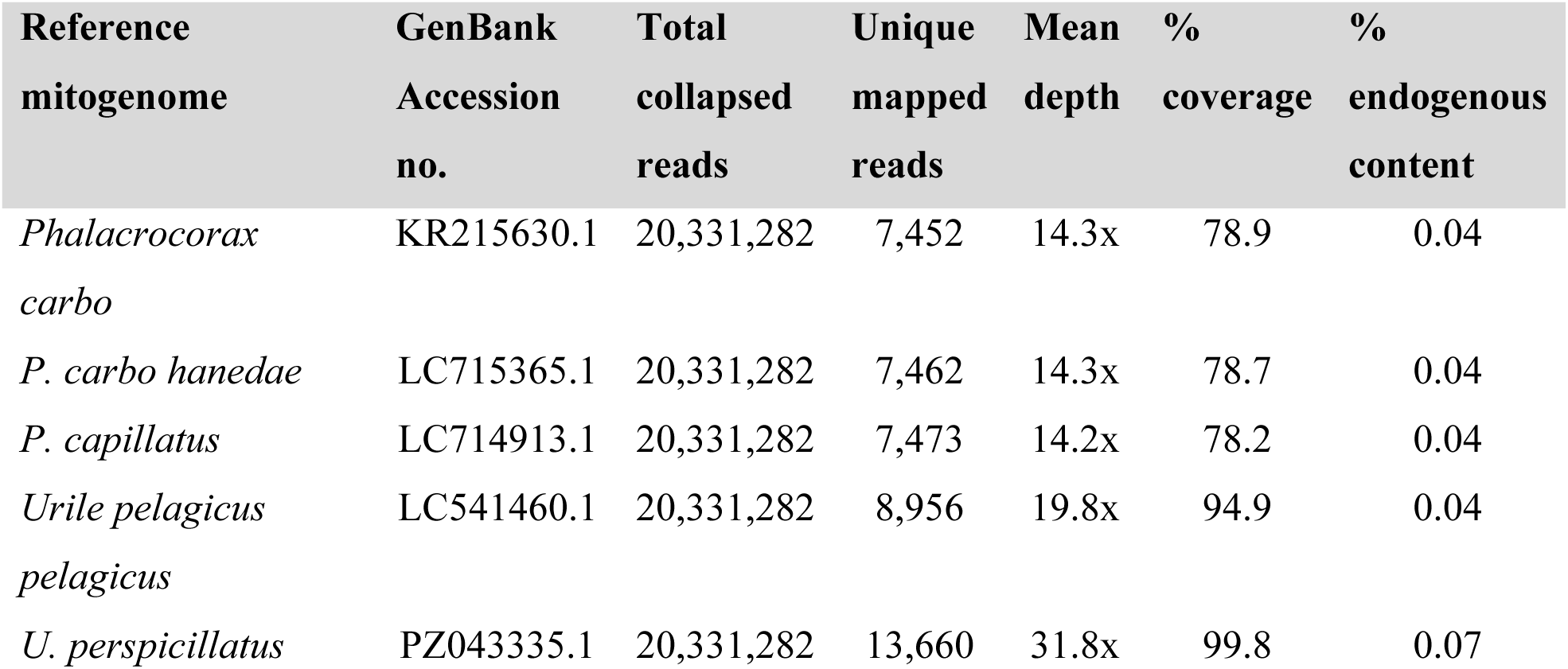
High-throughput sequencing and mapping statistics for selecting the optimal reference mitogenome for downstream mapping of Spectacled Cormorant (*Urile perspicillatus*) DNA sequences. The *U. perspicillatus* specimen (NHMUK 1842.12.21) with the highest endogenous DNA content from initial mapping to the Great Cormorant (*Phalacrocorax carbo*) reference nuclear genome was used for this comparison.

To assess the read distribution and depth of coverage across the five candidate reference mitogenomes, BAM files were imported into Geneious Prime v.2026.1.2 (https://www.geneious.com). The *U. perspicillatus* specimen (NHMUK 1842.12.21) with the highest endogenous DNA content from initial mapping to the *P. carbo* reference genome (GCA_963921805.1) was used for this comparison. The unpublished but recently publicly available *U. perspicillatus* (PZ043335.1) mitogenome, presumably from one of the historical skins in the Zoological Institute of the Russian Academy of Sciences, was subsequently selected to serve as the mitochondrial reference sequence as it resulted in the greatest number of mapped reads, highest depth of coverage, and highest endogenous content (Table 1). Consensus sequences for each *U. perspicillatus* sample were generated, calling nucleotides at bases with at least threefold coverage and 60% consensus, and assigning IUPAC ambiguity codes (N, R [A/G] or Y [C/T]) where coverage or consensus values fell below these thresholds. Consensus sequences were then aligned to the reference *U. perspicillatus* mitogenome using MAFFT v7.490 (Table 2; Katoh & Standley 2013), and annotations transferred using an 80% similarity threshold. Annotations and alignments were visually inspected for inconsistencies (e.g. internal stop codons), with none found.

**Table 2.** High-throughput sequencing and mapping statistics for all samples mapped against the Spectacled Cormorant (*Urile perspicillatus*) reference mitogenome.

| Sample ID | Total collapsed reads | Unique mapped reads | Mean depth | % coverage | % endogenous content |
| --- | --- | --- | --- | --- | --- |
| RMNH.AVES.107865 | 15,605,272 | 16,814 | 40.7x | 99.9 | 0.11 |
| KS.UL3639 | 12,010,427 | 31,411 | 107.1x | 99.9 | 0.26 |
| NHMUK 1842.12.21 | 20,331,282 | 13,660 | 31.8x | 99.8 | 0.07 |
| NHMUK 1858.2.31 | 13,278,281 | 31,010 | 105.9x | 99.9 | 0.23 |

Five mitochondrial genes (12S ribosomal RNA [12S], overlapping ATPase-8 and -6 [ATPase], NADH dehydrogenase subunit 2 [ND2], and cytochrome oxidase subunit I [COI]) were extracted and sequences were added to, and aligned with, a combination of the Kennedy *et al*. (2023), Rawlence *et al*. (2022), and Rawlence *et al*. (in review) datasets, including the nuclear FIB7, PARK7, IRF2, CRYAA, and RAPGEF1 genes (with additional Flightless Cormorant *Nannopterum harrisi* (Rothschild, 1898) sequences, which had previously been missing in those datasets, from Burga *et al*. (2017) [NEVG01000029]).

The models of nucleotide substitution for the Bayesian analysis were selected using the Akaike Information Criterion of Modeltest 3.7 (Posada & Crandall 1998). For this dataset the models selected for each gene region were: K81uf+I for 12S (6st+I), TrN+I for ATPase (6st+I), TIM+G for ND2 (6st+G), TVM+I+G for COI (6st+I+G), TIM+I for FIB7 (6st+I), HKY for PARK7 (2st), HKY for IRF2 (2st), K81uf+I for CRYAA (6st+I), and K81uf+I for RAPGEF1 (6st+I).

Bayesian phylogenetic analyses were performed using StarBEAST2 v1.0.0 (Ogilvie *et al*. 2017) as implemented in BEAST v2.7.7 (Bouckaert *et al*. 2019) to co-estimate the cormorant species tree together with the joint mitochondrial tree and the individual nuclear-gene trees. Overlapping ATPase-8 and -6 were concatenated, with this and other genes individually partitioned. We implemented an analytical population size integration model, unlinked substitution models for all partitions, linked trees for mitochondrial genes, and a birth-death species-tree prior. Uncorrelated lognormal relaxed molecular clocks were used because the phylogeny spans deep divergences among Pelecaniformes and Suliformes, over which substantial among-lineage rate variation may be expected; one linked clock for the nuclear genes and unlinked clocks for each of the mitochondrial genes.

The clock rates for the mitochondrial genes were modelled as normal priors with mean substitution-rates derived from rates estimated for terminal nodes 33–39 in Pacheco *et al*. (2011), corresponding to the region of the avian phylogeny encompassing cormorants and related seabirds, following Rawlence *et al*. (2022). The mean substitution rates in substitutions/site/million years (s/s/Ma) (and standard deviations) used were as follows: ND2: 0.00388 s/s/Ma (0.0013); COX1: 0.00232 s/s/Ma (0.0007); ATPase: 0.0029 s/s/Ma (0.0011); and 12S: 0.00145 s/s/Ma (0.0011). Mutation rates for the individual nuclear genes were modelled with uninformative 1/X priors.

The species tree was additionally constrained using three fossil calibrations modelled as lognormal priors. Following Prum *et al*. (2015), the Anhingidae–Phalacrocoracidae divergence was assigned a minimum age of 24.52 Mya based on the late Oligocene stem phalacrocoracid *Oligocorax stoeffelensis* (Mayr, 2007) (offset of 24.52 Mya, M = 4, and S = 0.8, mean in real space). A minimum age of 51.58 Mya, based on the early Eocene stem frigatebird *Limnofregata azygosternon* Olson, 1977, was used to constrain the deeper Pelecanus–Suliformes divergence (offset of 51.58 Mya, M = 9, and S = 0.6, mean in real space). Finally, the Morus–Sula divergence was constrained to a minimum age of 14 Mya (offset of 14 Mya, M = 6, and S = 0.6, mean in real space) based on a *Morus* sp. coracoid from the Langhian (16–14 Mya) of Portugal (Figueiredo *et al*. 2023).

We ran three independent MCMC chains, each run for 100 million steps, sampling every 10,000 steps. Convergence and sampling sufficiency were checked in Tracer v1.7.1 (Rambaut *et al*. 2018). For each independent MCMC run, we examined trace plots to confirm stationarity and mixing of all parameters. Effective sample sizes (ESS) were used to evaluate sampling sufficiency, with all parameters exceeding ESS values of 200 following removal of the 10% burn-in. Individual runs were combined in *LogCombiner* after discarding the first 10% of steps and maximum clade credibility consensus trees were generated in *TreeAnnotator* using the median node age.

## RESULTS

We obtained four complete *U. perspicillatus* mitogenomes (>99% coverage, average read-depth 31.8-107.1×; Table 2), which all exhibited patterns of nucleotide misincorporation consistent with post-mortem DNA damage (Supplementary Figure S1). All five *U. perspicillatus* mitogenomes (the four new ones and PZ043335.1) are identical, except for a single nucleotide polymorphism (SNP) in the control region of one individual, potentially indicative of a pronounced population genetic bottleneck.

Our Bayesian phylogenetic analysis (Figure 1B; Supplementary Figure S2) shows that *U. perspicillatus* is a strongly supported (posterior probability [PP] 1.0) monophyletic group within *Urile* cormorants (PP 1.0). There is moderate support (PP 0.90) for a sister relationship between *U. perspicillatus* and *U. pelagicus*/*U. urile* (Figure 1B), although we cannot rule out a sister relationship with the basal *U. penicillatus*.

Our molecular divergence dating (Figure 1B; Supplementary Figure S2) suggests *U. perspicillatus* diverged from *U. pelagicus*/*U. urile* 6.98 Mya (95% HPD 4.8-8.8 Mya). Variation within *U. perspicillatus* dates to 56 Kya (95% HPD 7-140 Kya), though given the single SNP amongst the five *U. perspicillatus* mitogenomes and the time-dependency of molecular rates (Ho & Larson 2006; Rawlence *et al*. 2022), this date may be an overestimate and should be treated with caution.

## DISCUSSION

Our phylogenetic analysis (Figure 1B; Supplementary Figure S2) and the osteological analysis of Samsonov *et al*. (2026) strongly supports that the Spectacled Cormorant belongs in the genus *Urile* and is most closely related to *U. pelagicus*/*U. urile*, although nuclear data is required to fully test this hypothesis given there is currently only mitochondrial sequence data for *U. perspicillatus* (e.g. Kennedy & Spencer 2014; Kennedy *et al*. 2019; Rawlence *et al*. 2022; Rawlence *et al*. in review).

The 6.98 Mya (95% HPD 4.8-8.8 Mya) divergence time between *U. perspicillatus* and its closest relatives suggests a long period of isolation. This depth of phylogenetic split is comparable to many of the deeper speciation events in Phalacrocoracidae, such as those between various *Phalacrocorax* species (Figure 1B; Supplementary Figure S2). In contrast, the equally high-latitude Southern Ocean *Leucocarbo* shags, except for the Rock Shag *L. magellanicus* (Gmelin, 1789), all diverged within the past 2.5 million years (Supplementary Figure S2; see also Rawlence *et al*. 2022).

The divergence of *U. perspicillatus* is broadly coincident with mid-late Miocene climatic cooling, diversification of marine kelp, and the origin of the modern kelp-forest ecosystem in the North Pacific during this time (Estes & Steinberg 1988; Estes *et al*. 2005; Starko *et al*. 2019). Highly productive kelp ecosystems have been hypothesized to have subsequently facilitated the diversification of large herbivorous invertebrates, benthic-foraging carnivores, and longer food chains (Vermeij 2012; Starko *et al*. 2019). We hypothesize that *U. perspicillatus* evolved its large size and benthic-foraging adaptations (see also Samsonov *et al*. 2026), to take advantage of this new niche, being able to handle rougher surf conditions, longer and deeper dives, and foraging for larger prey (Johnsgard 1993; Samsonov *et al*. 2026). While we cannot determine when *U. perspicillatus* evolved its large size, we hypothesize that it may have done so rapidly (e.g. Knapp *et al*. 2019; Verry *et al*. 2024; Tennyson *et al*. 2026).

The adaptations of *U. perspicillatus* may have facilitated ecological niche segregation with other sympatric but much smaller *Urile* cormorants in the coastal North Pacific. The early diverging North American *U*. *penicillatus* (7.9 Mya; Figure 1B) forages in inshore coastal zones and associated kelp-beds, utilizing the entire water column to hunt (Johnsgard 1993). In contrast, the late diverging North Pacific sister species *U. urile* and *U. pelagicus* (1.8 Mya; Figure 1B) are benthic foragers down to 30 m depth, predominantly of prey on rocky bottoms close to shore (Johnsgard 1993). While *U. urile* and *U. pelagicus* are morphologically similar in non-breeding plumage, these sister species are ecologically segregated through competitive exclusion of colony/nesting sites, with the larger *U. urile* potentially able to forage in deeper waters (Johnsgard 1993). The potential adaptations of *U. perspicillatus* may have also minimized competition with *P. carbo* (predominantly a benthic forager in waters less than 10 m deep) and *P. capillatus* (variable benthic and pelagic forager in waters up to 30 m deep) during the Pleistocene in Japan (Johnsgard 1993; Kato *et al*. 2002; Watanabe *et al*. 2018).

We argue that *U. perspicillatus* should be considered part of the North Pacific kelp-associated coastal fauna that evolved to inhabit these ecosystems, which were also distinguished by gigantic octopus *Enteroctopus dofleini* (Wulker, 1910), large molluscivorous fish (*Annarichthys* and *Bodianus* spp.), kelp-feeding sea cows (*Dusisiren* and *Hydrodamalis* spp.), sea lions (*Zalophus* and *Eumetopias* spp.), fur seals (*Callorhinus* spp.), tusked and tuskless walruses (*Dusignathus, Gomphotaria, Odobenus,* and *Valenictus* spp.), sea otters (*Enhydra* spp.), dwarf baleen whales (*Herpetocetus* spp.), flightless ducks (*Chendytes* spp.), and flightless auks (*Miomancalla* and *Mancalla* spp.; Miller *et al*. 1961; Domning 1978; Smith 2011; Vermeij 2012; Boessenecker 2013a; Boessenecker 2013b; El Adli *et al*. 2014; Estes *et al*. 2016; Boessenecker 2018; Vermeij *et al*. 2019; Boessenecker *et al*. 2024; Kiel *et al*. 2024). This fauna is unique in possessing the only cold-water herbivorous mammals and such a high diversity of vertebrates specialized for preying upon hard-shelled invertebrates (Vermeij 2012; Vermeij *et al*. 2019).

An interesting parallel with *U. perspicillatus* exists in the Pleistocene and Holocene of California, with the large flightless extinct duck *Chendytes lawi* Miller, 1925 (Howard 1955; Miller *et al*. 1961). *Chendytes* was an early diverging dabbling duck (Anatini) convergent with eiders (Mergini). This large-bodied (2-3.7 kg) species had an unusually robust skull adapted for feeding on marine invertebrates (perhaps for wrenching sessile species off rocky substrates), reduced forelimbs, and hindlimb adaptations for efficient deep diving (Miller 1925; Miller *et al*. 1961; Livezy 1993). Isotopic studies (Jones *et al*. 2021) confirm a diet rich in molluscs and fish, overlapping with sea otters *E. lutris* (Linnaeus, 1758) and harbor seals *Phoca vitulina* (Linnaeus, 1758). *Chendytes* persisted into the Holocene, being locally abundant in archaeological middens but declining gradually in representation until disappearing entirely about 2,500 years ago (Jones *et al*. 2008; Jones *et al*. 2021). It is tempting to speculate that *U. perspicillatus* may have been a boreal North Pacific analogue of *Chendytes*, with the evolution of both diving birds fostered by the high productivity and unusually diverse marine invertebrate assemblage of the North Pacific kelp-forest ecosystem. Isotopic analysis of fossil bones and historical skins of *U. perspicillatus* could shed further light on its foraging habits, especially in relation to other sympatric *Phalacrocorax* and *Urile* cormorants during the Pleistocene and Holocene.

We speculate that *U. perspicillatus* had a much wider prehistoric distribution around the North Pacific than is currently acknowledged, much like Steller’s sea cow (Domning 1978; Domning *et al*. 2007; Corbett *et al*. 2010; Balter 2012; West *et al*. 202l; Estes *et al*. 2015). The 120,000 year old *U. perspicillatus* bones from Shiriya Cape on northeastern Honshu, Japan, date to a period of increased ocean productivity that lasted until the Last Glacial Maximum (21,000 – 19,000 years ago; Watanabe *et al*. 2018). Given these bones were originally identified as *P. capillatus*, Holocene, Pleistocene, and even Pliocene fossil and archaeological assemblages from the coastal North Pacific, including those containing large cormorant bones, should be reassessed (e.g. Howard 1932; Howard 1949; Miller 1930; Miller 1971; Chandler 1990; Siegel-Causey *et al*. 1991; Lefevre & Siegel-Causey 1993; Lefevre *et al*. 1997), as misidentifications are common (e.g. Pleske 1896; Siegel-Causey *et al*. 1991; Olsen 2005; Watanabe *et al*. 2018; Mlikovsky 2023; Squires & Bond 2024; Samsonov *et al*. 2026). Re-examination of the fossil and archaeological record should also include bulk bone metabarcoding of unidentified ‘frag bags’ (e.g. Seersholm *et al*. 2018).

Our genetic analyses (Figure 1B; Supplementary Figure S2) suggest that *U. perspicillatus* either underwent a pronounced population bottleneck or exhibited long-term low effective population size, especially as it is likely only a single population was sampled given these cormorants were apparently restricted to Bering Island at the time of European discovery (Fuller 2000; Squires & Bond 2024). Given the extremely low genetic diversity within the five *U. perspicillatus* historical specimens, we suggest that a population bottleneck is more plausible. However, until palaeogenetic analyses are undertaken on the available late Holocene fossil remains (e.g. Squires & Bond 2024; Samsonov *et al*. 2026), we cannot determine whether this reduction in population size (1) represents an ancient event, post Last Glacial Maximum, associated with a range contraction to Bering Island as the climate warmed and ocean productivity changed (Watanabe *et al*. 2018), or (2) a more recent industrial scale human persecution. The impacts of 18^th^ and 19^th^ Century hunting and collecting practices could have also been exacerbated by millennia of subsistence hunting as humans moved through Beringia to settle the Americas ∼20,000 years ago (e.g. Moreno-Mayar *et al*. 2018) and, subsequently, the Aleutian Islands arc (i.e. islands that were not connected to the mainland at times of low sea stand) in a westward direction from 9,000 years ago to as recently as 2,500 – 500 years ago (Stanyukovich & Chernosvitov 1994; Corbett *et al*. 2010; Balter 2012; West *et al*. 2012).

It is likely that North Pacific kelp-associated ecosystems and the distribution of *U. perspicillatus* were heavily impacted by Pleistocene glacial cycles (Fraser *et al*. 2009; Watanabe *et al*. 2018) and human-induced trophic collapse (Estes *et al*. 2016), exemplified by the parallel range contractions of this taxon and the sympatric Steller’s sea cow (*H. gigas*) to an isolated relictual population. It has been hypothesized that the human-driven extirpation of *Enhydra* sea otters (in some areas after the decline of the large flightless duck *C. lawi*), a keystone species that predate on kelp-feeding sea urchins (Strongylocentrotidae), led to the trophic collapse of North Pacific kelp-ecosystems and subsequent contraction of the kelp-eating *H. gigas* to isolated and uninhabited islands where sea otters still occurred (Estes & Palmisano 1974; Estes *et al*. 2016; Jones *et al*. 2021). This scenario could explain the absence of both *U. perspicillatus* and *H. gigas* from archaeological deposits in the North Pacific. In addition, given that post-glacial sea level rise did not stabilize until ∼6,000 years ago, potential archaeological evidence of subsistence hunting of *U. perspicillatus* during the late Pleistocene and early Holocene may now be under water in some locations.

The speed of the over-exploitation and extinction of *U. perspicillatus* on Bering Island (ca. 100 years) may not have been enough time for a bottleneck to register in the mitogenome sequences of the individuals collected near the species extinction (e.g. Drummond *et al*. 2005 *cf.* Rawlence *et al*. 2015). Although we cannot confidently date the most recent common ancestor of the five *U. perspicillatus* mitogenomes due to their near-identical sequences, we suggest that the ancient bottleneck hypothesis may be more plausible and that the effective population size was likely quite small (despite being “exceedingly common” on Bering Island in 1741; Stejneger 1936).

Palaeogenomic analyses of *U. perspicillatus*, currently underway, will reconstruct the demographic history and genomic impacts of potential climate and (repeatedly) anthropogenically induced population bottlenecks and inbreeding on this enigmatic bird, testing the hypotheses raised in this study. Functional genomic analyses will also test for selection of SNPs associated with large body size, deep benthic foraging activity (in combination with dietary isotope analyses; Jones *et al*. 2021), and flight ability (e.g. Burga *et al*. 2017; Sackton *et al*. 2019; Edwards *et al*. 2024).

Historical museum specimens offer the only opportunity to interrogate the ecology and evolution of this now extinct and poorly known large seabird. Our analyses show that this enigmatic species had a long evolutionary history in the North Pacific and was likely an integral part of the kelp-associated fauna. The rapid extinction of *U. perspicillatus* highlights the magnitude of the impact humans have had on this highly productive coastal marine ecosystem.

## SUPPLEMENTARY DATA

Supplementary data are available at *Zoological Journal of the Linnean Society* online.

## ACKNOWLEDGEMENTS

We are grateful to the museum staff who supported destructive sampling of Spectacled Cormorant specimens: Mark Adams and Hein van Grouw (Natural History Museum Tring), Pepijn Kamminga (Naturalis Biodiversity Center, Leiden), and Alexander Alexio and Hannah Laakkonen (Luonnontieteellinen keskusmuseo, LUOMUS - Finnish Museum of Natural History, Helsinki). Thank you to Felix Marx and Charlie Boocock-Yee for helpful discussions about this manuscript.

## DATA AVAILABILITY

New mitogenome sequences generated as part of this study are available on GenBank (##XXXXXX-##XXXXXX), with raw sequencing reads available through FigShare: XXX.

---

**Supplementary Figure S1.**
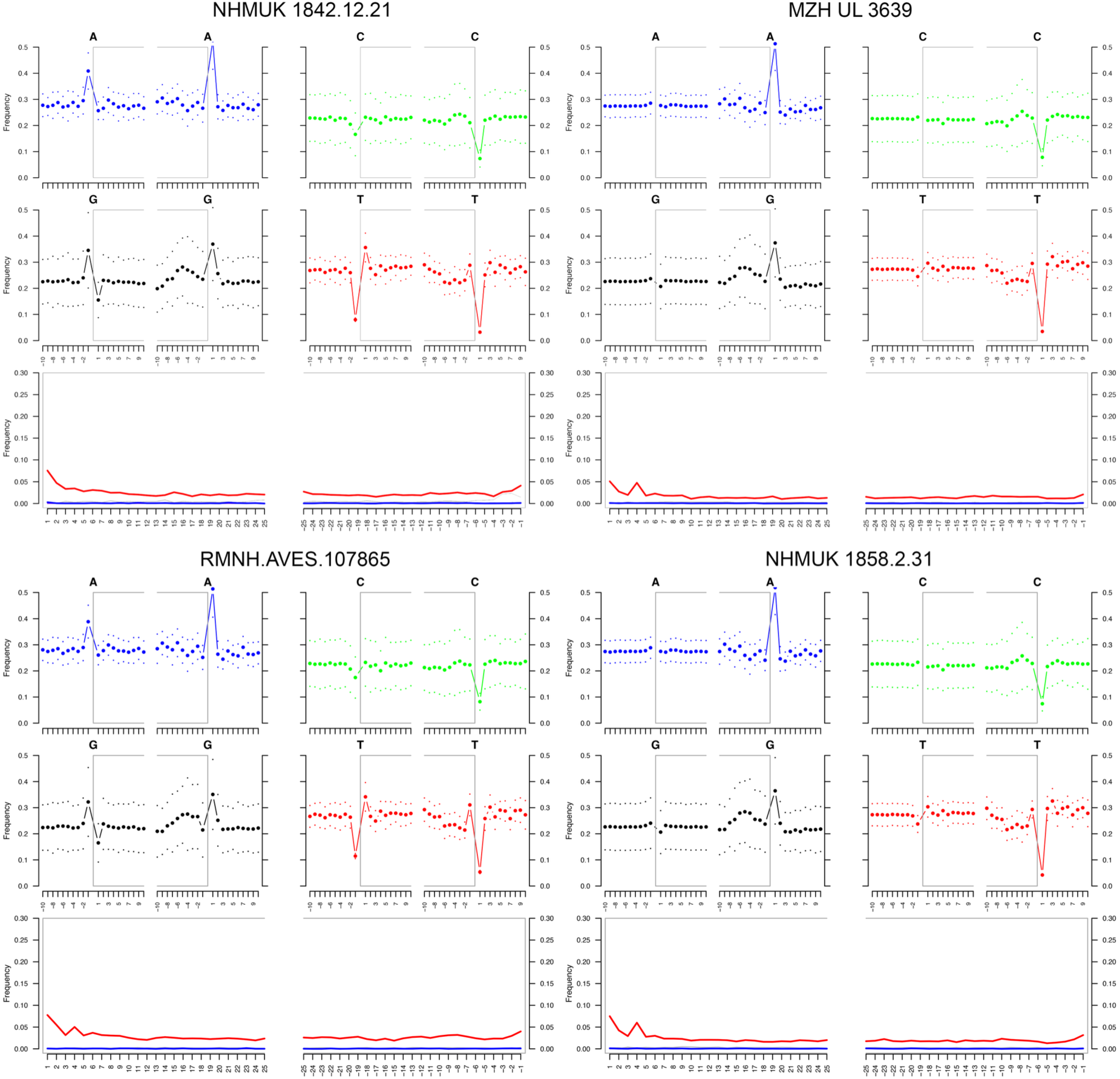
MapDamage plots for mapped collapsed reads of historic Spectacled Cormorant (*Urile perspicillatus*) single-stranded libraries.

**Supplementary Figure S2.**
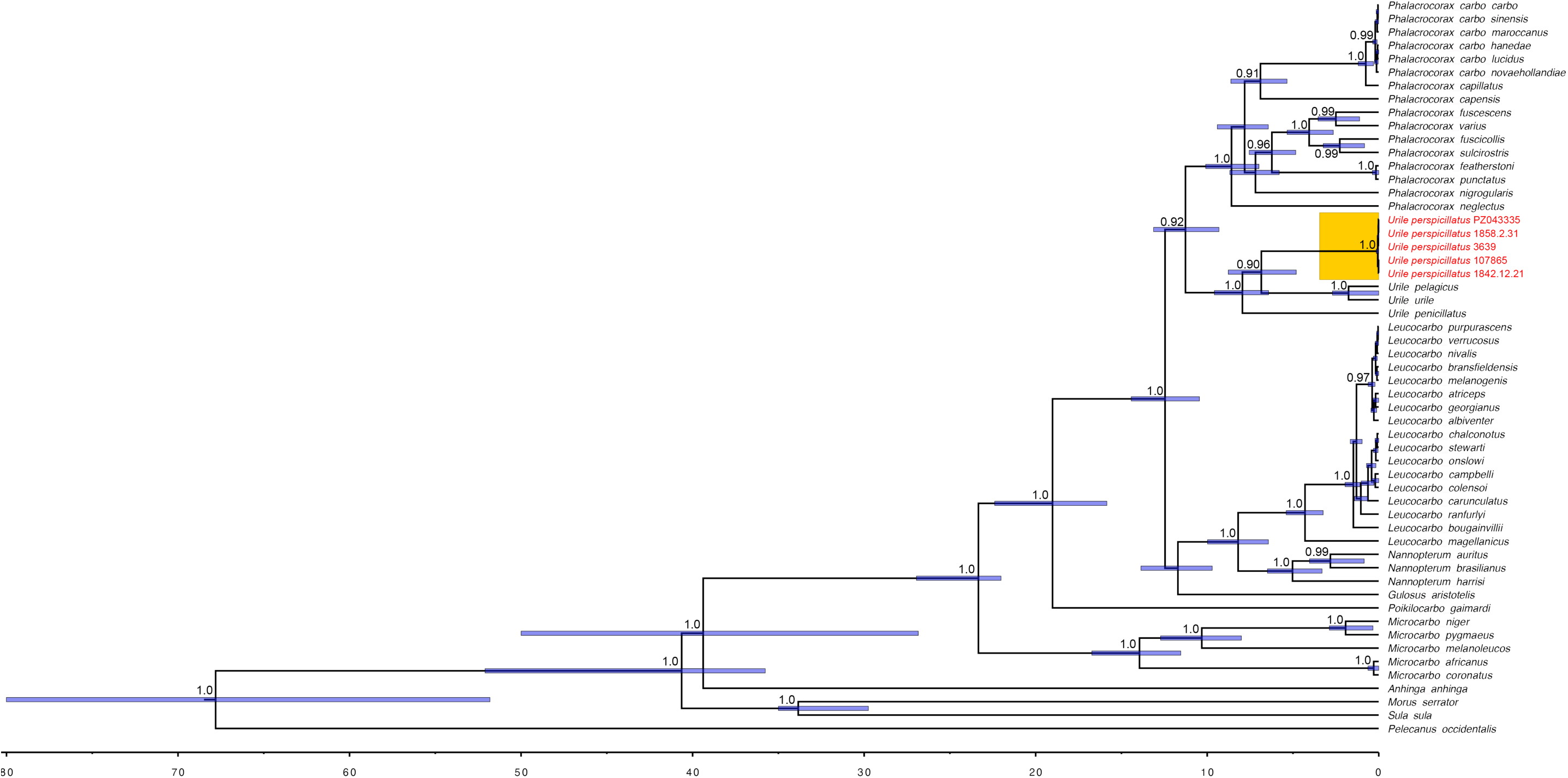
Evolutionary history of the Spectacled Cormorant (*U. perspicillatus*) and its nearest relatives. The time-calibrated phylogeny (in millions of years) is derived from 3.5 kilobases of mitochondrial DNA sequence data; the StarBEAST species-tree analysis additionally incorporated five nuclear loci for comparative taxa. Node bars on the phylogeny are 95% HPD of divergence times as indicated on the scale bar (millions of years before present). Node values are shown for BEAST Bayesian posterior probability support values of ≥0.90 for generic splits and those within *Phalacrocorax*.

